# Inconsistencies in *C. elegans* behavioural annotation

**DOI:** 10.1101/066787

**Authors:** Balázs Szigeti, Thomas Stone, Barbara Webb

## Abstract

High quality behavioural annotation is a key component to link genes to behaviour, yet relatively little attention has been paid to check the consistency of various automated methods and expert judgement. In this paper we investigate the consistency of annotation for the ‘Omega turn’ of *C. elegans*, which is a frequently used behavioural assay for this animal. First the output of four Omega detection algorithms are examined for the same data set, and shown to have relative low consistency, with F-scores around 0.5. Consistency of expert annotation is then analysed, based on an online survey combining two methods: participants judged a fixed set of predetermined clips; and an adaptive psychophysical procedure was used to estimate individual’s threshold for Omega turn detection. This survey also revealed a substantial lack of consistency in decisions and thresholds. Such inconsistency makes cross-publication comparison difficult and raises issues of reproducibility.

## 1 Introduction

Traditionally, behavioural annotation has been done manually, with the known weakness of inherent variability, as well as being labour intensive. In the current era of big data biology, there is an increasing tendency for behavioural annotation to be automated [1, 2]. Automated methods can obviously scale to significantly larger data sets, but they are also supposed to improve consistency by removing human judgement from the process. However, the self-consistency of automated methods does not guarantee consistency between different methods. Furthermore, these algorithms are typically validated relative to a human produced ‘ground truth’ dataset [3–7]. This evaluation process raises the possibility that algorithms are trained to learn the same observational biases - and variance - that are inherent to human annotation. Given that different research groups often use different annotation methods, a lack of consistency in their output could make comparison of published results from these groups difficult.

In this paper we specifically address the consistency of the behavioural annotation of the nematode worm *Caenorhabditis elegans* (*C. Elegans*), focusing on a particular worm behaviour, the Omega turn. Omega turns occur during reorientations, with the animal adopting a shape resembling the Greek letter Ω, see Figure 1**A** for a representative example. This behaviour was chosen as it is often treated as a discrete, well defined element of worm behaviour [5,7–10].

**Figure 1:**
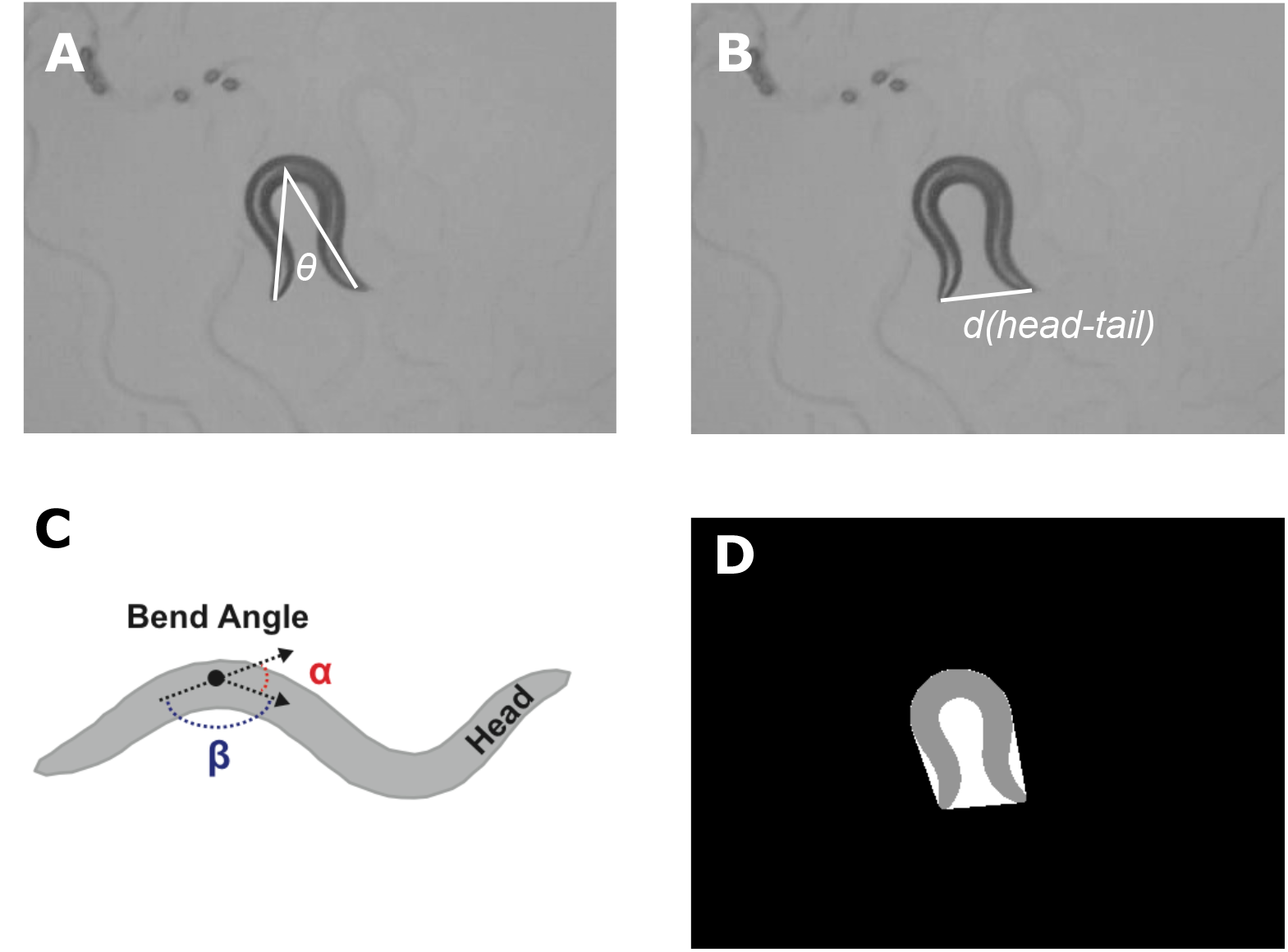
Visual explanation of the features that have been used construct the tightness score. Panel **A** shows the midbody angle *θ*, which is the angle between the head-middle and middle-tail vectors. Note that *π* – *θ* is the angle of reorientation of the event [6]. Panel **B** shows the head-tail distance. **C** illustrates worm bending that is measured using the supplementary angles to the bends formed along the skeleton. The bend angle (*α*) is the difference in tangent angles at each point; or, alternatively phrased, the supplementary angle (*α*) with respect to the angle formed by any three consecutive points (*β*). To detect Omega turns the midbody bend is calculated, which is the mean supplementary angle along the middle 1/3 of the worm’s body (image and caption is taken from [5]). Finally panel D introduces solidity, a measure of the overall concavity. It is defined as the ratio of the image (the worm’s body in grey) and the area of the convex hull (shown in white).

Our Omega turn consistency check has two components. First we examine the consistency of four Omega detection algorithms from the literature [4–7]. Second, we present the results of an on-line survey where we have invited experts to score Omega turns. The survey itself had two underlying components. Participants scored a set of predetermined clips and we have also employed an adaptive psychophysical method to identify individual’s threshold for Omega turns.

The results show that both expert annotation and algorithms are surprisingly inconsistent, and greater effort may be needed to ensure annotation methods provide a reliable basis for studies that include behavioural assays.

## 2 Methods

### 2.1 Behavioural data

This study used data from the *C. elegans* behavioural database (CBD) [5]. The database consists of worm videos and corresponding feature files that contain a number of precalculated feature time series (such as speed, eccentricity, eigenworm coefficients, etc.). We examined 776 experiments, all with hermaphrodite N2 worms. Worms were placed on a plate covered with a bacterial layer and the behaviour was recorded after a 30 minute habituation period. Each video is approximately 15 minutes long, so in total 194 hours of worm behaviour was analysed.

During Omega turns, the worm can contact itself, producing an intersecting shape in the videos, and for these frames it is difficult to extract a biologically meaningful skeleton [6, 11]. As a consequence these ‘coiling’ frames are not processed in the CBD and the features for the corresponding frames are not calculated. If the resulting gap in the video was smaller than 20 consecutive frames (0.6 sec) then linear interpolation was used to gain a proxy for the features. This interpolation method is not reliable for longer gaps, hence Omega events that contained longer gaps were discarded.

### 2.2 Consistency of Omega turn detection algorithms

#### Algorithms

Four algorithms have been taken from the literature to examine their consistency with each other. The algorithms are from the Zentracker package [4], the *C. elegans* behavioural database (CBD) [5], a computer vision based study to detect such events [6] and from a recent publication studying search behaviour [7]. Common to all these methods is that they detect Omega turns if a feature or a combination of features exceeds a user defined threshold. For example, [5] uses the midbody bend as the defining property of Omega turns. Note that this is not an exhaustive list of Omega turn detection algorithms. These particular algorithms have been chosen because the code used for the original publication was readily available.

#### Consistency quantification

To summarise annotation consistency we report the precision (positive predictive value) and sensitivity (also known as recall and true positive rate) [12]. Precision is the ratio of true positive events to all events recognised, while sensitivity is the proportion of true positives to all reference events. Mathematically they are expressed as

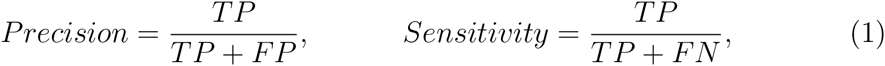

where *TP, FP* and *FN* are true positive, false positive and false negative respectively. For example, if one algorithm is taken as the reference for Omega events, a true positive occurs for the comparator algorithm when it selects the same event (a *TP* was counted if at least 50% of the frames identified as part of an Omega turn overlapped); a false positive when it selects an event not labelled by the reference algorithm; and a false negative when it fails to select an event that was labelled by the reference algorithm. Precision and sensitivity are often combined to a single number summary, the F-score, which is defined as:

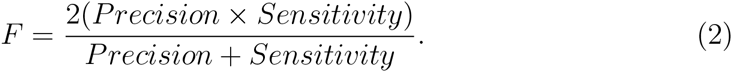

#### Threshold tuning

The consistency between annotation algorithms is likely to be affected by parameter settings. Therefore we calculated the results first with the original feature thresholds (taken from the publication) for each method, and then with the thresholds altered so as to find the best match between each pair of algorithms that could be obtained by parameter adjustment.

To find the best match, each algorithm was run 25 times with different thresholds. For each run the difference in the threshold was increased or decreased by 2.5% of the initial value. Therefore a range 70%-130% of the initial threshold values were scanned. Lower percentages correspond to a more permissive definition (i.e. more events classified as Omega turns), but some scales had to be inverted. For example [4]’s method uses an upper bound on ‘eccentricity’ and a lower bound on ‘solidity’. Therefore to make the run associated with 70% more permissive, the eccentricity scale had to be inverted.

### 2.3 Community survey of Omega turns

#### Survey structure

To compare the consistency of expert Omega turn detection an online survey was developed^1^. After a brief registration, participants were shown 40 short (2-5s) clips of Omega events and were asked to indicate, using a button press, if each was an Omega turn or not. Participants were also asked to rate their confidence to detect Omega turns on a scale 1-5 (with 5 being very confident).

In the survey we wanted to include ambiguous, wide amplitude turns that one may or may not consider an Omega turn. Therefore to select events for the survey we have run the Omega detection algorithm by [6] on the CBD videos, but with the threshold reduced to 75% of its original value. Using this criteria 1526 Omega like events were detected.

The 40 clips in the survey were made up of two components. There was a set of 20 predetermined videos that were scored by everyone. The remaining 20 were determined by an adaptive threshold finding procedure, where the next clip shown depends on previous answers. Specifically, the truncated staircase method was used [13] to estimate thresholds (see below). To conceal this structure and reduce order effects these two components (predetermined set and threshold finding) have been mixed together such that each predetermined clip was followed by a clip used to detect the threshold. The participants were not told in advance of these two underlying components to eliminate possible cognitive biases.

To gather responses to the survey, we emailed 47 experts (PIs identified from publications on *C. elegans* behaviour) inviting them and their laboratory members to participate. The survey was also advertised through the social media presence of the OpenWorm project.

#### Selection of predetermined clips

To select the 20 predetermined clips, the eigenshape annotator (ESA) was used [3]. In brief ESA is an unsupervised behavioural annotator that produces a probabilistic annotation. Events were selected that are labeled as Omega turns, but had a high entropy (0.75*H_max_* ≤H), i.e. Omega events were selected that had a high classification uncertainty. 158 events met this criteria and from this set 20 were selected randomly, see the online *Supplementary videos* to watch the clips.

#### Adaptive threshold finding

To deploy an adaptive threshold finding technique, it was necessary to have a single metric by which Omega turns could be ranked. We developed a ‘tightness’ metric score based on the Omega turn detection algorithms in the literature. Most Omega turn detection algorithms recognise such events when a certain feature exceeds a user defined threshold. Features that are commonly associated with Omega turns are solidity, midbody angle, head-tail distance and midbody bend. For a visual explanation for each of these features see Figure 1.

For each Omega event the peak amplitude of these features were measured. Across all events the z-score was calculated for each feature peak and the tightness score of each event is the mean z-score across the four features. This procedure ranks the Omega-like events from wide amplitude turns to the sharper, more ‘characteristic’ Omega turns. It is not claimed that the tightness score captures every variation of Omega like events. However it quantifies the sharpness of coils that is the key feature of turning behaviours. For a demonstration of the resulting ranking see the online *Supplemental Video 1*.

A truncated staircase method was used to estimate an expert’s omega detection threshold (measured on tightness score) [13]. The equation to select the next clip is

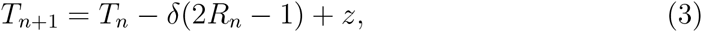

where *δ* is a fixed step size (in tightness score), *T_n_* is the tightness of the clip shown at the *n^th^* step and *R_n_* is the *n^th^* response (*R_n_* = 1 is the answer is yes and *R_n_* = 0 if the answer is no) and *z* is a small random variation to avoid repetitions. In this process the sequence of clips has either increasing or decreasing *T_n_* until a switch in the subject’s response (from yes to no, or no to yes) for successive clips occurs. In this case the step direction is reversed and again the stimulus strength (*T_n_*) monotonically increases or decreases until the next switch in response. To estimate the threshold, the average *T_n_* at the points where the subject switched responses is taken.

## 3 Results

### 3.1 Consistency of Omega detection algorithms

The consistency of four Omega turn detection algorithms was quantified. In Table 1 the precision, sensitivity and F-score of the methods are presented relative to each other. The scores are calculated first using the prameter settings originally provided, and then when the parameters of two methods were tuned for optimal match in outputs (results given in brackets; for details of the tuning procedure see the *Threshold tuning* section). Without tuning, the results show little consistency, with an average F-score of 0.3. Even with tuning to find the best match, the F-score frequently stays below 0.5, indicating poor consistency in classification.

**Table 1:** Consistency of Omega turn detection algorithms. The top of each column shows which algorithm was taken as reference and the rows correspond to the algorithm being compared to it. In each cell the *Precision/Sensitivity/F – score* are reported, for a description of these measures see the section *Consistency quantification*. The numbers in parentheses in each cell report the same statistics with thresholds tuned for optimal match, see the section *Threshold tuning* for further details.

|  | Huang 2006 | Yemini 2013 | Salvador 2014 | Laurent 2015 |
| --- | --- | --- | --- | --- |
| Huang 2006 | 1/1/1 | 0.40/0.46/0.43<br>(0.64/0.65/0.65) | 0.28/0.15/0.20<br>(0.52/0.38/0.43) | 0.13/0.67/0.22<br>(0.79/0.69/0.74) |
| Yemini 2013 | 0.46/0.4/0.42<br>(0.66/0.67/0.67) | 1/1/1 | 0.45/0.22/0.29<br>(0.66/0.43/0.51) | 0.05/0.21/0.08<br>(0.92/0.69/0.79) |
| Salvador 2014 | 0.15/0.28/0.20<br>(0.48/0.52/0.43) | 0.26/0.5/0.34<br>(0.47/0.71/0.56) | 1/1/1 | 0.12/0.1/0.11<br>(0.62/0.83/0.77) |
| Laurent 2015 | 0.68/0.13/0.22<br>(0.64/0.79/0.74) | 0.22/0.05/0.1<br>(0.7/0.93/0.8) | 0.62/0.1/0.13<br>(0.83/0.72/0.77) | 1/1/1 |

The Omega detection threshold was also estimated for each algorithm using the same methodology as for expert annotations (see *Adaptive threshold finding*). The results are shown on Figure 2B, for this figure the original parameters from the publications were used. Note that in agreement with Table 1 there is overlap in the confidence intervals, but there is no clear consensual threshold.

**Figure 2:**
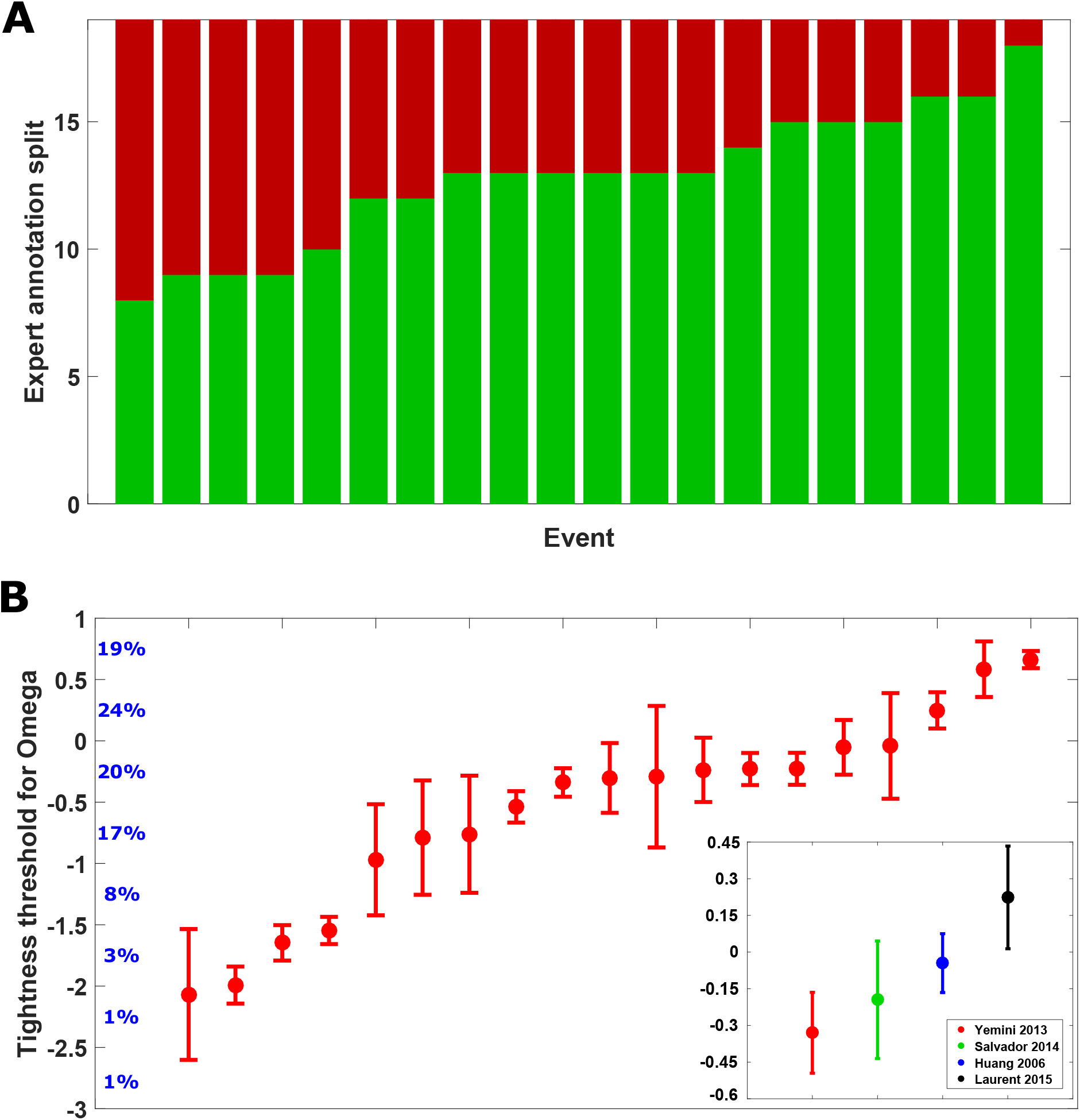
Outcomes of the Omega turn community survey. The data was filtered to exclude non-expert annotations, see the *Consistency of expert annotation* for details. Top panel shows the split of experts (green: ‘yes, it was an Omega’; red: ‘not an Omega’) for the set of 20 predetermined clips, ordered by the proportion of experts who agreed it was an Omega turn, which ranged from 8/19 to 18/19. Panel **B** shows the results of the threshold determination procedure. Each data point is one expert’s estimated tightness threshold to detect Omegas with the corresponding 95% confidence interval, ordered by increasing tightness. Inset shows the estimated threshold and confidence intervals for the Omega detection algorithms. Blue numbers next to the y-axis indicate what percentage of the data (all potential Omega events, see *Survey structure*) falls between tightness z-scores (e.g. 19% of the events had a tightness z-score between 0.5 and 1). This shows wide divergence in how many events different experts would classify as an Omega turn.

The algorithm by [4] produces the worst match to the other algorithms. This is due to the method only picking out the sharpest of Omega turns, hence it identifies many fewer events compared to the other methods. It is not argued that any of the methods assessed is worse or better than the others, but rather the point is that results could differ significantly depending on which method a particular analysis uses.

### 3.2 Consistency of expert annotation

Overall 27 survey responses were collected in the period 2016 May 30 - June 14. For the results presented here we have discarded the responses whose confidence in detecting Omega turns was below 4, so only expert annotation is analysed (19 participants in total).

As described in the *Methods*, the survey had two components: a set of predetermined clips and an adaptive threshold finding procedure. Figure 2**A** shows the distribution of answers for the predetermined clips, which had been selected for high classification uncertainty according to an unsupervised behavioural annotator (see *Methods*). None received a unanimous consensus, and only 6 were judged the same by more than 75% (at least 15 out of 19) of the experts. Almost half the clips produced a split of 12:7 or worse.

The estimated decision thresholds for each expert and the corresponding 95% confidence intervals are shown on Figure 2**B**. Note the different size of confidence intervals reflects the number of samples to estimate the threshold, which depends on the number of switch points from yes to no for each subject in the sequence of 20 presentations (see *Adaptive threshold finding*). It is nevertheless also an indicator of the subject’s (internal) consistency as more switch points, and hence smaller C.I., suggests the staircase quickly converged to oscillate around a specific value. It is clear that the estimated thresholds spread widely, with no region where the majority cluster, or all confidence intervals overlap.

## 4 Discussion

In this paper we have shown that both automated and expert annotations of *C. elegans*’s Omega turns are surprisingly divergent. First the implications for worm research are discussed. Then some general comments regarding supervised behavioural analysis is presented. Finally we speculate whether the observed annotation inconsistency is a more general feature of behavioural studies.

Characterising *C. elegans* behaviour often involves an estimate of Omega turn probability [3–7,9]. It is important to check whether the algorithms used to detect Omega turns are consistent, otherwise it is difficult to make cross-publication comparisons. It was found that the four Omega turn detection algorithms we tested produce a surprisingly divergent annotation even after their respective parameters have been adjusted for optimal match.

One way to overcome the inconsistency problem would be if the community adopted the same platform for behavioural analysis. There is a range of publicly available packages [3–5], however, each comes with its own strengths and weaknesses, hence it is difficult to see the whole community adopting any one of these methods. A potential solution would be an open-source software that is developed and maintained not by a single laboratory, but rather by the whole research community. This way each lab would have ownership and the cross talk between laboratories could lead to a deeper appreciation of the limitations of each analysis technique.

A potential source of the inconsistency we have observed is that the Omega turn is not a distinct behaviour, but rather a part of a spectrum of turning behaviours. We have previously argued for this possibility based on the high proportion of uncertain classification of behavioural events [3]. Others have also supported this hypothesis based on the geometry of locomotion states [14] and based on the continuous neuronal representation of motor sequences [15].

A major limitation of our work, in both our earlier paper and the current publication, is that events could not be analysed where the worm was intersecting itself for an extended period (see *Methods*). Recently a method was developed that can resolve coiling postures [11]. Their analysis of eigenworm amplitudes found a multi-modal distribution that could be used as a data driven definition of Omega turns. Furthermore this study reports that ‘beyond’ Omega turns there is another sharper turning behaviour, the Delta turn.

However one should note that in this study the experimental conditions were not identical to ours. In the CBD data (used here) worms are browsing in food, while in this study worms were analysed off food. The 1^*st*^ and 3^*rd*^ eigenworms switch position (sorted by eigenvalues) in these two conditions indicating that the behaviour is altered (off food the first two eigenworms are associated with locomotion and the 3^*rd*^ one is associated with turns, on food the 1^*st*^ eigenworm corresponds to the turning postures) [5, 16]. Therefore the results may or may not generalise to other experimental conditions.

Our analysis of expert annotation has general implications for supervised approaches to behavioural analysis. The common element to these methods is that they take an investigator labeled dataset and then an algorithm learns to reproduce the expert annotation [1]. As a consequence, supervised methods can be only as consistent as their training data. Therefore prior to using supervised methods we would urge investigators to first examine the variability of expert opinion. Furthermore we note that unsupervised methods are often evaluated against a human produced ‘ground truth’ dataset. This evaluation process imposes subjective factors and hence leads to similar problems as with the supervised methods. The validation of unsupervised methods is a complex issue that raises many philosophical questions as well [17, 18].

Although we have only analysed one specific behaviour of one model organism, the observed inconsistencies in behavioural annotations (both expert and automated) seem likely to be more widespread. For example there is an analogous uncertainty about how to define the behavioural states of larval *Drosophila melanogaster* [3,19–22]. Different publications use different ways of defining the behavioural states, most likely due to the difficulty in finding an unambiguous characterisation. As a result, a similar inconsistency of the various analysis techniques should be a cause for concern in reproducibility of maggot research. We hope that with our analysis we have inspired investigators to carefully look at the issue of consistency for other model organisms as well.

## Acknowledgements

The authors would like to express their gratitude for Aidan Rocke for his initial work on this project. Furthermore we would like to thank Andre Brown, Emanuel Busch and members of the Insect Robotics group for their feedback on the survey prototype. This work was supported by grants EP/F500385/1 and BB/F529254/1 for the University of Edinburgh School of Informatics Doctoral Training Centre in Neuroinformatics and Computational neuroscience from the UK Engineering and Physical Sciences Research Council (EPRSC), UK Biotechnology and Biological Sciences Research Council (BBSRC), and the UK Medical Research Council (MRC), and the FP7 program MINIMAL.

## Author contributions

BS conceived the study, developed the code, analysed the data and wrote the article. TS developed the web implementation of the Omega event selection algorithm and maintained the survey’s website. BW has supervised the project and helped to write the manuscript.

## Footnotes

1 The survey can be reached at http://groups.inf.ed.ac.uk/worms/index.html

